# Plasmonic Nanocavity-Based Method for Measuring Electric Charges in Solution

**DOI:** 10.1101/2025.03.19.644097

**Authors:** Alexey I. Chizhik, Damir I. Sakhapov, Jörg Enderlein, Narain Karedla

## Abstract

Electric charges play a fundamental role in shaping the structure, function, and interactions of biomolecules, yet precisely measuring these charges at the single-molecule level remains a significant technical challenge. Here, we introduce a novel experimental methodology that utilizes plasmonic nanocavities to quantify molecular electric charges in solution with high sensitivity. Our approach exploits an externally applied electric field to induce the spatial redistribution of charged molecules confined within a planar metallic nanocavity, while simultaneously leveraging nanocavity-induced fluorescence lifetime modulation as a highly sensitive readout. We demonstrate the feasibility of this method through proof-of-concept experiments, where we measure the fluorescence lifetimes of positively and negatively charged fluorescent dye molecules as a function of the applied electric field across the cavity. The experimental results are validated through a rigorous theoretical framework, incorporating statistical thermodynamics and electrodynamic modeling to accurately describe the observed data. The proposed method offers a calibration-free, experimentally simple, and rapid alternative to existing charge measurement techniques, opening new avenues for precise quantification of molecular electric charges down to the single-molecule level.

## Introduction

Electric charges play a fundamental role in the structure, function, and interactions of biomolecules, governing a wide range of biological processes essential for life. Biomolecules such as proteins, nucleic acids, and lipids carry charged functional groups that are in equilibrium with the ions in solution and with each other, influencing their stability, dynamics, and interactions with their environment [1–9]. Understanding these electrostatic properties is crucial for deciphering molecular mechanisms in biology, from enzymatic catalysis to cellular signaling, as well as for understanding protein folding and aggregation.

However, precisely measuring the electric charges of molecules in solution remains a challenging task, particularly at low molecular concentrations or even at the single-molecule level. State-of-the-art optical microscopy-based techniques, such as the anti-Brownian electrokinetic trap and evanescent scattering microscopy, have been explored for quantifying molecular charges at the single-molecule level [10, 11]. However, these approaches are either technically challenging and restricted to one single molecule at a time or require molecular modification and tethering to a surface, limiting their broader adoption.

Recently, Krishnan et al. introduced an exciting new approach for single-molecule charge measurements by exploiting electrostatic trapping in micro-wells and analyzing the escape probability of trapped molecules [12–16]. The core idea involves trapping freely diffusing molecules within nanoscale indentations etched into glass microchannels, which possess a high surface electric charge. These surface charges generate a repulsive electrostatic potential for like-charged molecules in solution, with a local energy minimum near the center of the indentation that transiently confines them. By measuring the Brownian motion-driven average escape time of a molecule from the indentation, its electric charge can be inferred: the longer it remains trapped, the larger its charge.

Although these approaches are highly innovative, they suffer from low throughput, limited flexibility, and considerable experimental complexity. To obtain quantitative charge values, the surface charge of the nanoindentations should be precisely known, and complex calculations are required to determine the three-dimensional electric field distribution. Additionally, modeling Brownian motion and escape probability within the intricate indentation geometry adds further computational challenges.

Here, we propose a novel approach for measuring molecular electric charges in solution by integrating two core principles: (i) utilizing an external electric field to induce the redistribution of charged molecules within a nanocavity-enclosed solution, and (ii) employing nanocavity-induced fluorescence quenching as a readout for this redistribution. By monitoring the redistribution of charged molecules as a function of the externally applied electric field, we can directly measure the electric charges of the enclosed molecules. Furthermore, as we demonstrate in this paper, this approach also enables the simultaneous determination of the ionic strength of the solution.

The schematic of the plasmonic nanocavity setup is shown in Fig. 1. A solution containing the fluorescently labeled molecule of interest is enclosed within a planar nanocavity of sub-wavelength dimensions. In our experiment, the nanocavity is formed by a planar coverslip and a concave glass lens with a sufficiently large curvature radius. This design ensures that when the fluorescence excitation laser is focused into a diffraction-limited spot within the cavity, close to the lens’ symmetry axis, the illuminated cavity volume can be approximated as a nearly plane-parallel cavity. This plasmonic nanocavity design has been extensively used in the past for quantum yield measurements of luminescent emitters [17–20].

**FIG. 1.**
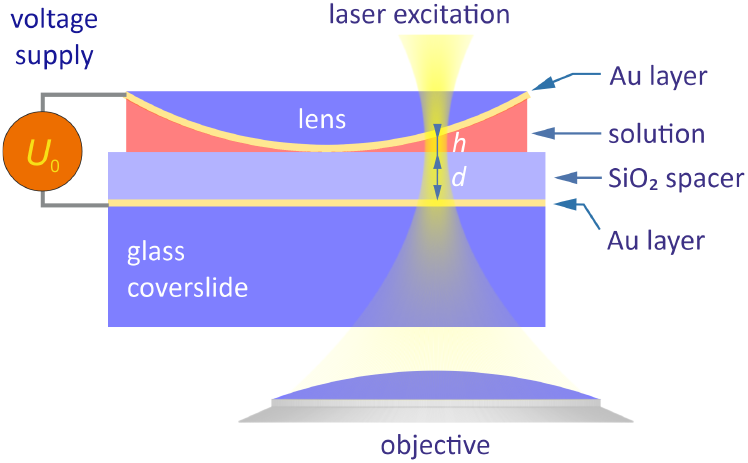
Schematic of the plasmonic nanocavity. For details see main text.

On both sides, the cavity is enclosed by two thin gold layers, which serve a dual purpose. Firstly, applying a voltage to these layers generates a non-uniform electric field across the cavity. Secondly, the gold layers induce position-dependent fluorescence quenching: the farther a fluorescent molecule is from the surfaces, the greater its fluorescence emission brightness and fluorescence lifetime. This effect arises from the near-field coupling of the fluorophore’s excited state to free electron oscillations (plasmons) in the metal layers. This coupling accelerates the transition from the excited state to the ground state, thereby shortening the fluorescence lifetime. Additionally, due to partial dissipation of the excited-state energy into surface plasmons, fluorescence brightness diminishes as the molecule approaches the metal surface.

In this paper, we focus exclusively on the effect of fluorescence lifetime reduction, which is inherently calibration-free and independent of molecular concentration, making it a more robust and reliable measurement compared to fluorescence brightness analysis.

### Theory

Firstly, we consider the non-uniform distribution of charged molecules within the nanocavity that arises when applying a voltage difference between the two gold electrodes. This applied voltage induces a nonuniform electric potential distribution across the dielectric SiO_2_ spacer layer of thickness *d* and the solution layer of thickness *h*.

The electric potential in the SiO_2_ layer (−*d < x <* 0) follows the Laplace equation:

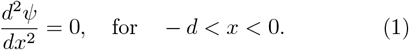

In contrast, within the solution layer (0 *< x < h*), the potential is governed by the linearized Poisson-Boltzmann equation [21]:

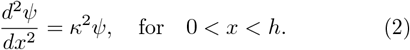

Here, *κ* is the inverse Debye length, defined as:

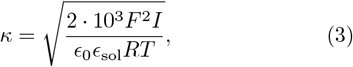

where *F* and *R* are the Faraday constant and the universal gas constant, respectively. The parameters *ϵ*_0_ and *ϵ*_sol_ denote the dielectric constants of vacuum and the cavity-enclosed solution, respectively, while *T* represents the temperature. The ionic strength *I* (in M) is given by:

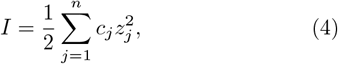

where *c*_*j*_ and *z*_*j*_ correspond to the molar concentrations and charge numbers of all *n* ionic species present in the solution.

For an applied voltage difference *U*_0_, the boundary conditions for the electric potential are:

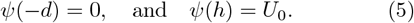

Additionally, at the SiO_2_/solution interface (*x* = 0), the continuity conditions

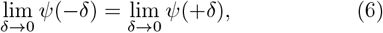

and

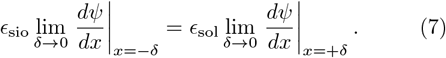

must hold. The solution to these equations is given by:

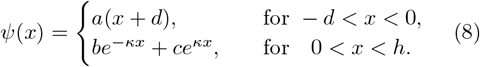

where the constants *a, b*, and *c* are determined from the boundary and continuity conditions.

With the electric potential distribution *ψ*(*x*) known, the spatially varying concentration *c*_*j*_(*x*) of the *j*^th^ species of charged molecules follows the Gibbs-Boltzmann distribution:

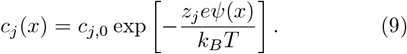

Figure 2(a) presents the numerical results for the electric potential, while Figure 2(b) illustrates the corresponding molecular concentration distributions. The calculations were performed using the following parameters: an applied voltage of 1 V, a *d* = 100 nm-thick SiO_2_ spacer layer with a dielectric constant of *ϵ*_sio_ = 3.7, and a *h* = 100 nm-thick solution layer with a dielectric constant of *ϵ*_sol_ = 80. The solution contains only two oppositely charged monovalent ionic species (*z*_*j*_ = ± 1), each at an equal concentration of 10 µM, resulting in an ionic strength of *I* = 10 µM. For comparison, the uniform distribution of a non-charged species is also depicted (green line).

**FIG. 2.**
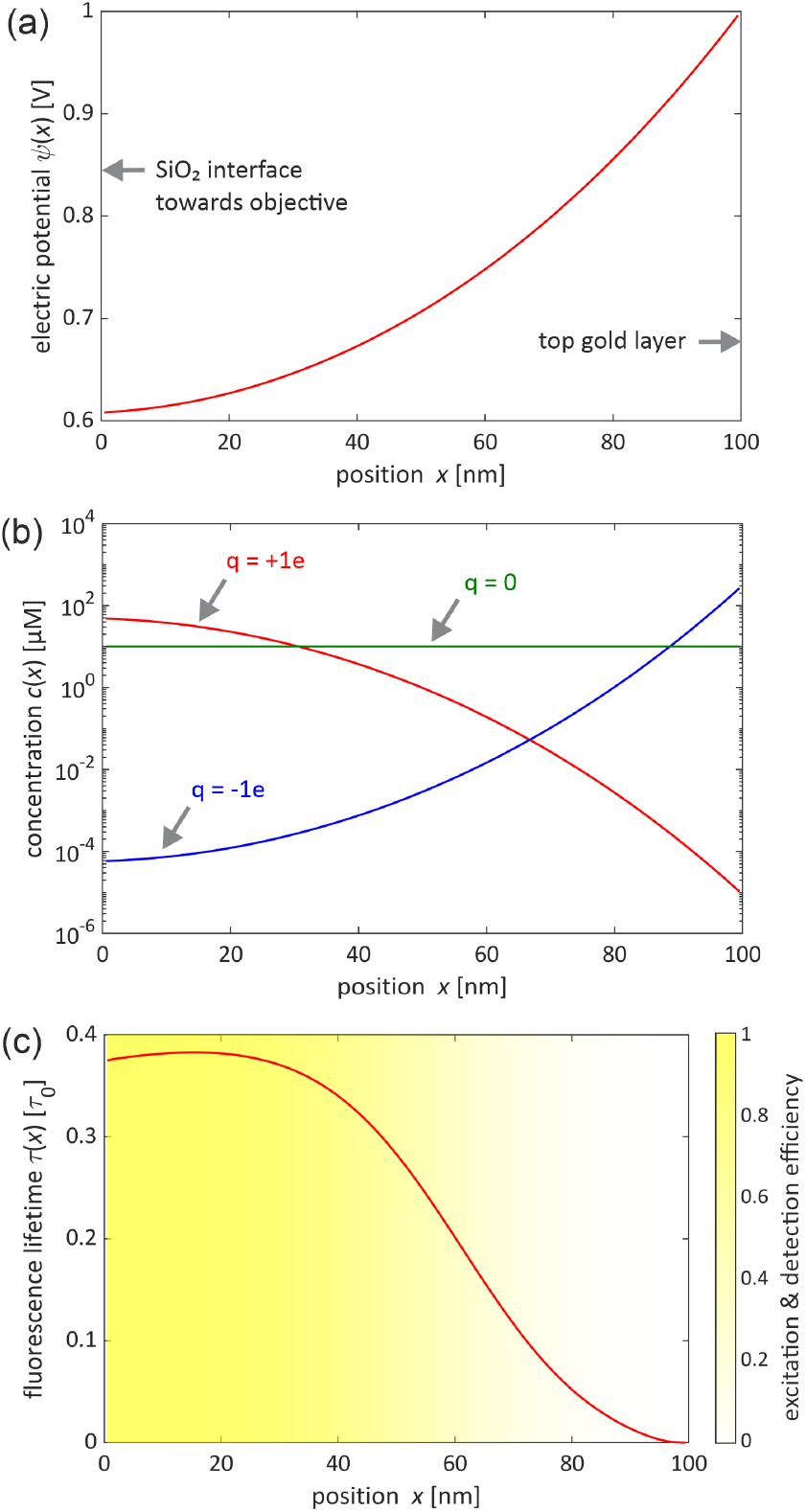
Panel (a) displays the electric potential across the solution-filled cavity, panel (b) shows the resulting concentration distributions, and panel (c) illustrates the position-dependent fluorescence lifetime. In panel (c), the shading and color bar represent the position-dependent detection efficiency (a.u.).

Secondly, it is necessary to model the mean fluorescence lifetime measured within the detection volume, which is defined by the intersection of the fluid-filled nanocavity and the diffraction-limited excitation focus. The excitation intensity generated by the focused laser light within the solution is calculated by representing the electric field of the Gaussian excitation beam as a plane wave expansion and then using Fresnel’s equations for plane wave reflection/transmission for calculating the resulting electric field distribution (and thus intensity) within the cavity, see also [17].

The modification of a molecule’s excited-state lifetime due to its coupling with cavity modes—primarily governed by near-field interactions with free electrons in the metal layers—can be modeled by treating the emitting molecule as an ideal electric dipole emitter [22]. By solving Maxwell’s equations for the given nanocavity, one can determine the position- and orientation-dependent emission rates of the dipole. This approach has been extensively detailed in refs. [17, 23] in the context of tunable metallic nanocavities used for modulating single-molecule emission and measuring quantum yields. Here, we briefly summarize the main concepts.

It is assumed that the fluorescing molecules are sufficiently small such that their rotational diffusion time in solution is much shorter than the fluorescence decay time—a condition that is well satisfied by small fluorophores used in this study (with typical rotational diffusion times of a few dozen picoseconds compared to fluorescence lifetimes of several nanoseconds). Under this assumption, the orientation-averaged emission rate is given by:

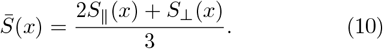

where *S*_∥,⊥_(*x*) are the emission rates for horizontally (⊥, perpendicular to the *x*-axis) and vertically (∥, parallel to the *x*-axis) oriented dipoles, which depend only on the dipole’s position along the *x*-axis.

Given the fluorescence quantum yield *ϕ* of the dye, and considering that the emission rate *S*_0_ of a unit-amplitude dipole emitter in an unbounded homogeneous medium with refractive index *n*_sol_ is *S*_0_ = *c*^2^*n*_sol_*/*3 (in cgs units), where *c* is the vacuum speed of light, the fluorescence lifetime *τ* (*x*) of a fluorophore at position *x* (i.e., the inverse of its emission rate) is:

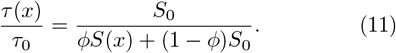

where *τ*_0_ is the fluorescence lifetime value that would be measured for molecules in bulk solution without any nanocavity enclosure. Figure 2(c) shows the dependence of the fluorescence lifetime on the position in the nanocavity in units of *τ*_0_. The calculations were done for two gold layers (top and bottom) of 15 nm thickness with an additional 1 nm titanium adhesion layer between the gold and the glass/SiO_2_ interfaces. The calculations were done for an emission wavelength of 570 nm, and the corresponding values of the complex refractive indices for the metals were taken from ref. [24]. As evident from the figure, the presence of the metallic nanocavity induces a strong fluorescence lifetime gradient across the solution gap, enabling the detection of even subtle concentration shifts of fluorescent molecules within the solution.

To obtain the mean fluorescence lifetime that would be measured experimentally, one also needs to account for the position-dependent detection probability *p*_det_(*x*), which describes the likelihood of detecting a photon from a molecule at position *x*. This probability is determined by integrating the dipole’s orientation-averaged angular emission distribution over the objective’s cone of light collection and normalizing it by the total dipole emission emission rate, as obtained by solving Maxwell’s equations. This position-dependent probability alias detection efficiency is shown as a density plot underlying the lifetime curve in Figure 2(c). Finally, the observable mean fluorescence lifetime is computed as:

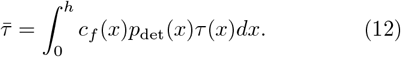

where *c*_*f*_ denotes here the non-uniform concentration distribution of the fluorescent molecules within the non-uniform electrostatic potential, see Figure 2(b).

### Experiment

For proof of principle, we measured pure solutions of two different dyes: the singly positively charged dye Atto Rho6G amine (*τ*_0_ = 4.1 ns, *ϕ* = 0.94) and the singly negatively charged dye Atto 488 azide (*τ*_0_ = 4.1 ns, *ϕ* = 0.8), under neutral pH conditions. Experimentally, we employed the plasmonic nanocavity shown in Figure 1 in combination with a custom-built scanning confocal microscope equipped with a fluorescence lifetime imaging extension. The cavity mirrors were fabricated by vapor deposition of gold onto the surface of a clean glass coverslip (bottom mirror, thickness 170 µm) and a plane-convex lens (top mirror, focal distance 150 mm) using a Leybold Univex 350 evaporation system under high-vacuum conditions (10^−6^ mbar). The bottom and top gold films had a thickness of 15 nm. To reduce fluorescence quenching by the bottom metal film and to provide electrical insulation between the metal mirrors, the bottom mirror was further coated with a 100 nm-thick SiO_2_ layer. For improved adhesion of the gold layers to the glass substrates and the SiO_2_ layer to the gold film, additional 1 nm thick titanium layers were deposited. Electric cables were attached to the gold films using a conductive epoxy (Chemtronics, CW2400), and the voltage was applied using an electric power supply unit (Thorlabs, BPC303).

Fluorescence lifetime measurements were performed with a high-numerical-aperture objective lens (Apo N, 60×, 1.49 NA oil immersion, Olympus). The excitation source was a pulsed white-light laser system (NKT Photonics, WL-MICRO), and fluorescence detection was achieved using a single-photon detection module (Micro

Photon Devices, PDM series photon counting detector module). Data acquisition was carried out with a multichannel picosecond event timer (HydraHarp 400, PicoQuant). Fluorescence decay curves were obtained by generating histograms of photon arrival times with a bin width of 32 ps.

Measurements were performed at the optimal cavity length, where the brightest fluorescence was observed. This condition corresponds to the condition where the maximum emission spectra of the molecules are in resonance with the cavity mode. This results in solution gaps of 86 nm for Atto 488 azide (emission maximum 520 nm) and 100 nm for Atto Rho6G amine (emission maximum 557 nm). All fluorescence decay curves were recorded until reaching 10^4^ counts at the maximum. The fluorescence decays, *F* (*t*), were truncated at 70 ps after the excitation pulse peak and fitted using a multi-exponential decay model, from which the average excited-state life-time was calculated as:

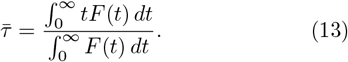

Mean fluorescence lifetime values were measured for various voltage amplitudes, ranging from − 1 V to 1 V, applied sequentially at both electrodes.

Since the results are highly sensitive to the ionic strength of the solution, we initially used a dye concentration with known charge in the micromolar range and then fitted the ionic strength value using our model calculations. Taking this as a reference, we repeated the measurements for three solutions with ionic strengths of 1 µM, 10 µM, and 100 µM. Figure 3 presents the experimental results for both molecules: the positively charged amine of the fluorescent dye Atto Rho6G and the negatively charged azide of the fluorescent dye Atto 488.

**FIG. 3.**
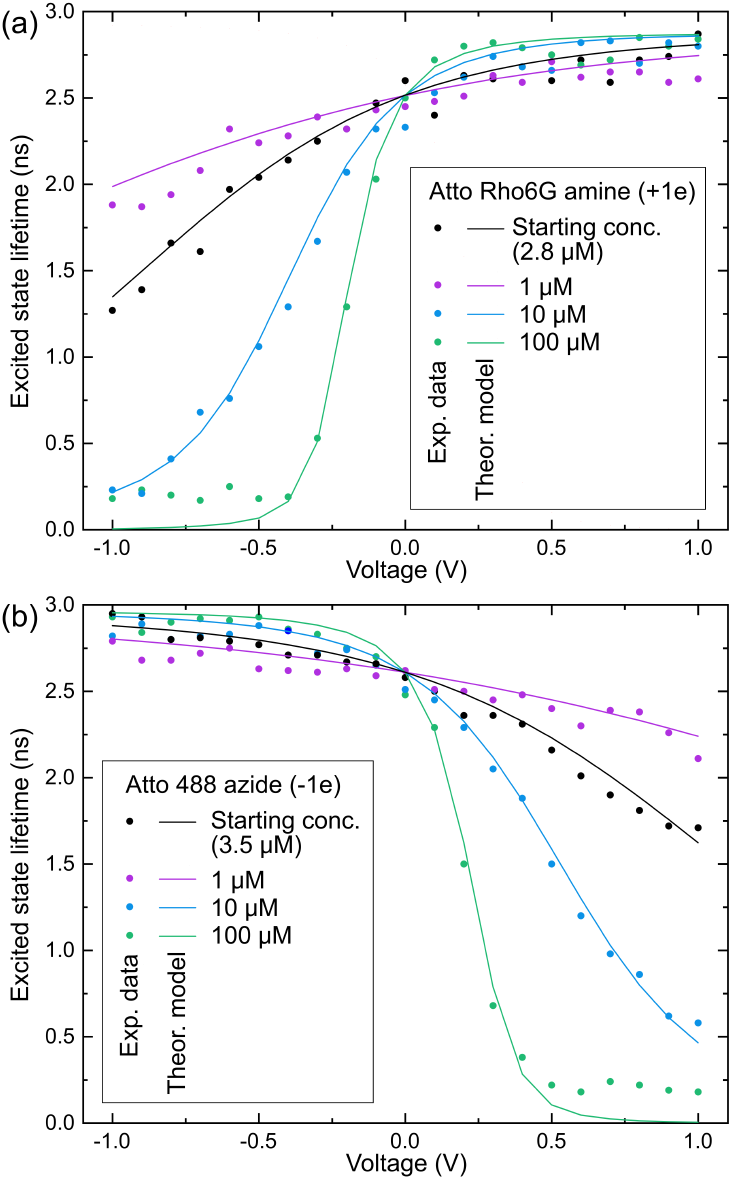
Experimental results. Shown are the measured fluorescence lifetime from dye solutions as a function of applied voltage and for solutions of different dye concentration alias ionic strength. Circles are measured lifetime values, solid lines fits using the theoretical framework explained before. Panel (a) shows the result for pure solutions of Atto Rho6G-amine in water (electric charge +1 e), and panel (b) a similar result for the dye Atto488-azide (electric charge −1 e).

As shown in Figure 3, the theoretical model provides a strong fit to the experimental curves. It is important to emphasize that this close agreement between theoretical predictions and experimental data is only achievable when the electric charge of the dyes is precisely ± 1 e, as assumed. Any deviation from this charge value would result in increasing discrepancies between the shape and position of the theoretical curves relative to the experimental data. Furthermore, as expected, we observe a complete reversal of the lifetime-versus-voltage curves when the sign of the electric charge is switched, further validating the model’s predictive accuracy. The noticeable scattering of the measured lifetime values around the theoretical values is *not* due to measurement errors in the fluorescence lifetime, as these are consistently smaller than the size of the circular markers in the plots. Instead, we attribute the observed scattering to imperfect screening of external electric fields and the limited accuracy of the voltage supply, which should be improved in future experiments. More interestingly, at higher ionic strengths (∼ 100 µM) and larger absolute voltages (ca. ± 1 V), we observe a systematic deviation in which the experimental lifetime values exceed the theoretical predictions. This discrepancy clearly arises from the breakdown of the validity of the *linearized* Poisson-Boltzmann equation. To achieve greater accuracy, future work should employ the *nonlinear Poisson-Boltzmann equation*, despite the significantly higher computational complexity of its numerical solution.

## Conclusion

The primary objective of this study was to introduce a novel methodology for the challenging task of measuring the electric charge of molecules in ionic solutions and to demonstrate the experimental feasibility of the plasmonic nanocavity approach. As a final remark, it is worth noting that our method can be extended to measure charges of single molecules or to analyze a solution containing a mixture of molecules with different charges. This could be achieved by employing an extended planar nanocavity combined with a high-sensitivity fluorescence-lifetime wide-field detection scheme (see e.g. ref. [25]), enabling single-molecule fluorescence measurements over sufficiently long time periods (e.g., via single-molecule tracking).

## ACKNOWLEDGMENTS

J.E. acknowledges financial support by the European Research Council (ERC) for financial support via project “smMIET” (grant agreement no. 884488) under the European Union’s Horizon 2020 research and innovation program. A.I.C. is grateful to Ingo Gregor and Oleksii Nevskyi for technical support.

